# Mechanical Suppression of Breast Cancer Cell Invasion and Paracrine Signaling Requires Nucleo-Cytoskeletal Connectivity

**DOI:** 10.1101/838359

**Authors:** Xin Yi, Laura E. Wright, Gabriel M. Pagnotti, Gunes Uzer, Katherine M. Powell, Joseph Wallace, Uma Sankar, Clinton T. Rubin, Khalid Mohammad, Theresa A. Guise, William R. Thompson

**Affiliations:** Department of Physical Therapy, School of Health and Rehabilitation Sciences, Indiana University, Indianapolis, IN 46202; Department of Medicine, Division of Endocrinology, School of Medicine, Indiana University, Indianapolis, IN 46202; Department of Mechanical and Biomedical Engineering, Boise State University, Boise, ID 83725; Department of Biomedical Engineering, Purdue School of Engineering and Technology, Purdue University, Indianapolis, IN 46202; Department of Anatomy & Cell Biology, Indiana University, Indianapolis, IN 46202; Department of Biomedical Engineering, Stony Brook University, Stony Brook, NY11794

**Keywords:** LINC, MDA-MB-231, Vibration, Osteolysis, Osteoclastogenesis, Nesprin, Sun

## Abstract

Exercise benefits the musculoskeletal system and reduces the effects of cancer. The beneficial effects of exercise are multifactorial, where metabolic changes and tissue adaptation influence outcomes. Mechanical signals, a principal component of exercise, are anabolic to the musculoskeletal system and restrict cancer progression. We examined the mechanisms through which cancer cells sense and respond to mechanical signals. Low-magnitude, high-frequency signals were applied to human breast cancer cells in the form of low-intensity vibration (LIV). LIV decreased invasion through matrix and impaired secretion of osteolytic factors PTHLH, IL-11, and RANKL. Furthermore, paracrine signals from mechanically stimulated cancer cells, reduced osteoclast differentiation resorptive capacity. Physically disconnecting the nucleus by knockdown of *SUN1* and *SUN2* impaired the ability of LIV to suppress invasion and production of osteolytic factors. LIV also increased cell stiffness; an effect dependent on an intact LINC complex. These data show that mechanical signals alter the metastatic potential of human breast cancer cells, where the nucleus serves as a mechanosensory apparatus to alter cell structure and intercellular signaling.

## Introduction

Physical activity has beneficial effects on nearly every organ system. In addition to the positive effects of exercise on cardiovascular(Sattelmair et al., 2011) and musculoskeletal health(Warden and Thompson, 2017), regular physical activity is associated with a reduced risk of colon, endometrial, and breast cancers(Moore et al., 2016; Thune and Furberg, 2001). Women who exercise, at a moderate intensity for 3-4 hours per week, have a 30-40% reduced risk of breast cancer, compared to sedentary women(Friedenreich and Orenstein, 2002). Additionally, physical activity is associated with decreased cancer mortality(Li et al., 2016) and reduced tumor size in mice(Pedersen et al., 2016), even at low doses(Zhao et al., 2019). While these studies highlight the positive effects of exercise on cancer, the underlying mechanisms remain largely unknown.

Exercise inherently involves repetitive bouts of physical movement, altering whole-body homeostasis with subsequent adaptations at the cell, tissue, and organ levels(Warden and Thompson, 2017). The beneficial effects of physical activity on cancer seems, at least partially, due to metabolic and immune effects. Exercise results in reduced insulin resistance and decreased hyperinsulinemia in muscle(Frank et al., 2005). Additionally voluntary running results in reduced tumor size in mice, due to increased recruitment and infiltration of natural killer (NK) immune cells(Pedersen et al., 2016), suggesting that exercise regulates cancer growth partially through improved immune responses. While the effects of exercise on cancer are multifactorial, the contribution of mechanical force, a principal component of physical activity, in regulating cancer progression is unclear.

While exercise suppresses tumor growth and reduces cancer-related mortality, musculoskeletal complications arising from cancer treatments and cancer itself make exercise difficult at best, or physically dangerous at worst. The subsequent sedentary state perpetuates the problem, as the mechanical signals imparted through exercise are absent. Prior work demonstrates that the physical signals necessary to activate cellular responses need not be large, nor of long duration(Thompson et al., 2014; Zhao et al., 2019). As such, introducing very low magnitude mechanical signals exogenously may provide the necessary benefits of mechanical input while avoiding the negative consequences of more strenuous forms of traditional exercise.

Low magnitude mechanical forces can be introduced to the musculoskeletal system through platforms that emit low intensity vibration (LIV) signals, serving as an effective “exercise surrogate” by delivering mechanical input similar to that of exercise(Rubin et al., 2001). When applied to mesenchymal progenitor cells LIV promotes proliferation and differentiation(Pongkitwitoon et al., 2016). At the molecular level, LIV signals initiate a signaling cascade resulting in increased phosphorylation of focal adhesion kinase (FAK) and Akt, resulting in downstream activation of RhoA and formation of filamentous actin structures(Thompson et al., 2013). The effects of LIV are additive, with a second bout of LIV enhancing FAK phosphorylation and F-actin contractility(Uzer et al., 2015).

Previous work in non-cancerous cells demonstrated that the anabolic effects of mechanical stimuli are enhanced when a refractory period is introduced between loading bouts(Sen et al., 2011). As such, the responses produced with two 20-minute bouts separated by three hours were greater than that of a single 40-minute mechanical stimulus. Similar effects were observed in animal models(Patel et al., 2017). These data suggest that low magnitude signals are sufficient for anabolism, but also that proper dosing primes the cells to generate a more robust response with subsequent mechanical stimuli.

The mechanical compliance of tumor cells dictates cell behavior, where stiffness of the plasma membrane is inversely proportional to metastatic potential(Mohammadi and Sahai, 2018). Cells with decreased stiffness display increased migration and invasion, which is regulated by the organization of the actin cytoskeleton(Zhou et al., 2017). Exogenous mechanical input enhances actin cytoskeletal structure(Thompson et al., 2013), and work in non-cancerous cells demonstrates that the nucleus serves as a critical mechanosensory organ where direct connections between the nucleus and the cytoskeleton enable transmission of low magnitude, oscillatory mechanical signals(Uzer et al., 2015). Attachment of the nucleus to the cytoskeleton is enabled by the LINC complex, containing Nesprin and Sun proteins, and may be a means by which exercise influences the metastatic properties of cancer cells. Further, cells from human breast tumors have decreased expression of Nesprin and SUN(Matsumoto et al., 2015), suggesting that the LINC complex may regulate tumorigenicity. In this work, we subjected human breast cancer cells to mechanical signals and examined direct biochemical changes of the cancer cells, indirect paracrine signaling alterations, and the biophysical mechanisms enabling transmission of mechanical signals to breast cancer cells.

## Results

### LIV does not directly alter cell death but increases the susceptibility to Fas ligand-induced apoptosis

Mechanical signals regulate cell death of several cancer types(Lien et al., 2013). To determine if direct application of LIV to breast cancer cells influences cell death, MDA-MB-231 human breast cancer cells were exposed to twenty-minute bouts of LIV (0.3 g, 90 Hz) once- or twice-daily for three days, in the presence or absence of TGF-β1. Control cells were placed on the vibration platform with no transmission of LIV signal. Cell viability was assessed using the 3-(4,5-Dimethylthiazol-2-yl)-2,5-Diphenyltetrazolium Bromide (MTT) assay. No changes in cell viability were observed with LIV or TGF-β1 treatment (Fig. 1a). mRNA expression of *FAS*, a cell membrane death receptor, was measured by quantitative polymerase chain reaction (qPCR). Once-daily LIV increased *FAS* expression by 2-fold, but this change was not significant (Fig. 1b). Expression of *FAS* was significantly (p<0.05) increased by 3-fold following twice-daily LIV (Fig. 1b). Expression of *CD95*, the gene encoding for Fas ligand was not altered with LIV (Fig. 1c).

**Figure 1.**
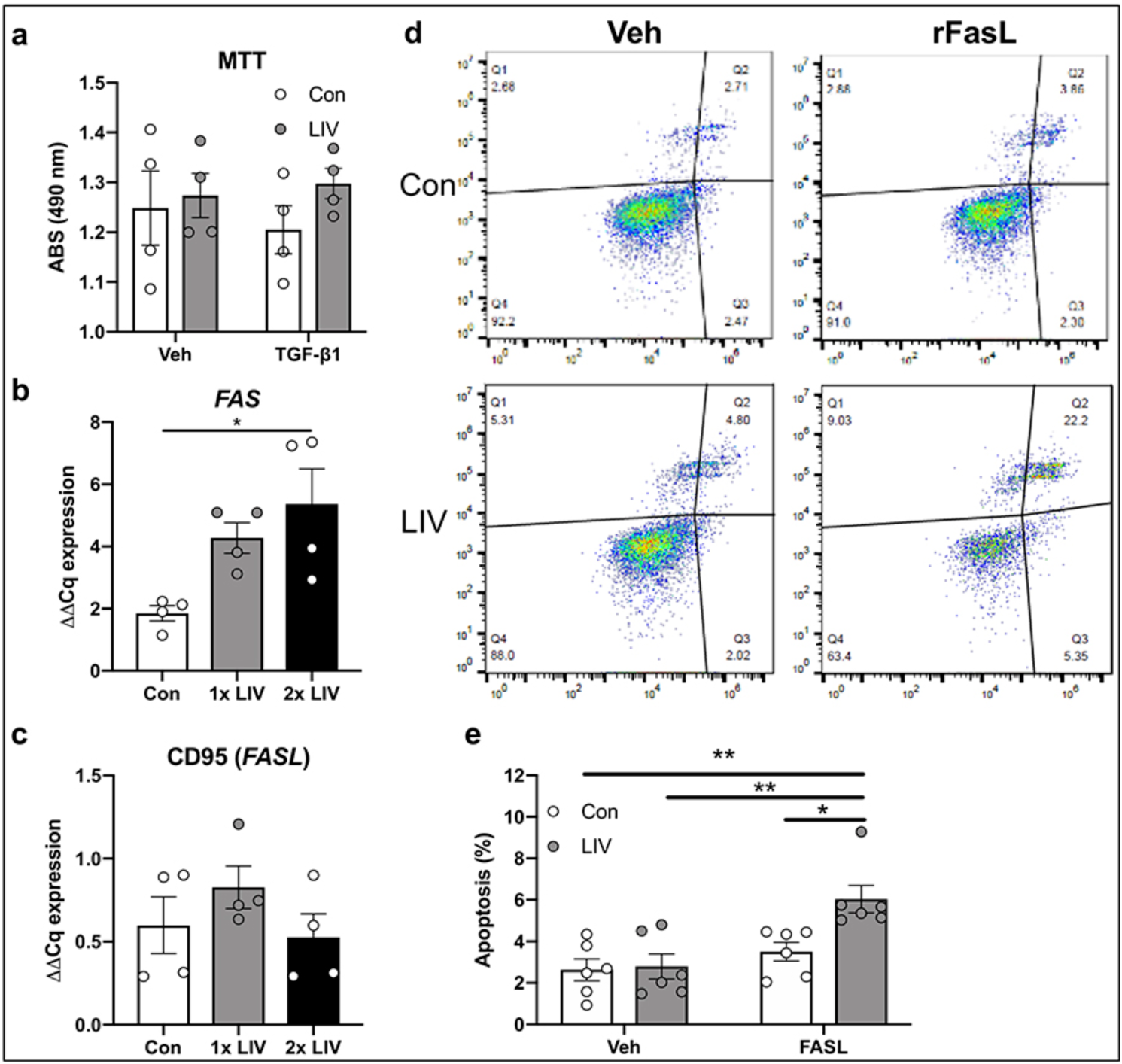
LIV increases susceptibility of FasL-mediated apoptosis. (a) MTT assay demonstrated no changes in cell viability, in the presence of absence of TGF-β1 following twice-daily LIV (2x LIV). Points represent individual measurements of biological replicates (n=4). (b) Expression of *FAS* was quantified by qPCR and normalized to *GAPDH* (n=4). Twice-daily LIV (2x LIV) significantly increased *FAS* expression, no significant difference was found with once-daily LIV (1x LIV). (c) qPCR analysis of *CD95*, normalized to *GAPDH* (n=4). (d) Images obtained from flow cytometry of Annexin V-stained MDA-MB-231 cells. Quadrant #3 (Q3) represents early apoptosis. (e) Quantification of flow cytometry data from quadrant #3 following once- or twice-daily LIV, and treatment with PBS (veh) or recombinant Fas ligand (FASL). Data represents six independent biological replicates. One-way ANOVA (b & c) or Two-way ANOVA (a & e) p-values: *p<0.05, **p<0.01.

LIV-induced upregulation of *FAS* led us to hypothesize that expression of the Fas death receptor may increase the susceptibility of MDA-MB-231 cells to Fas ligand-mediated apoptosis. As such, MDA-MB-231 cells were exposed to twice-daily LIV or placed on LIV platforms with no signal transmitted (control) and treated with recombinant Fas ligand (75 ng/ml) for 24 hours prior to staining with Annexin V and subsequent sorting by flow cytometry (Fig. 1d). No changes in apoptosis were observed following LIV in vehicle-treated cells (Fig. 1e). Treatment with Fas ligand did not alter apoptosis in control cells; however, addition of Fas ligand to cells treated with LIV increased (p<0.05) the percent of apoptotic cells compared to control cells by ∼2-fold (Fig. 1e). These data demonstrate that application of LIV does not directly induce cell death, but that MDA-MB-231 cells are more susceptible to Fas ligand-mediated apoptosis following exposure to mechanical signals.

### Low magnitude mechanical signals suppress invasion

Extravasation and subsequent metastasis of cancer cells requires invasion through matrix-dense borders. To determine if exogenously applied LIV influenced the ability of MDA-MB-231 cells to invade through ECM, trans-well invasion assays were performed. Cells were treated with LIV for twenty minutes per bout, once- or twice-daily for three days. Cells then were trypsinized and seeded onto trans-well membranes containing Matrigel^®^ and visualized using crystal violet (Fig. 2a). Once-daily LIV induced a 1.4-fold reduction in invasion; however, cells exposed to LIV twice-daily had a significant (p<0.01) 3-fold reduction (Fig. 2b). The area of cells that invaded through the trans-well membrane was also quantified, showing a significant (p<0.05) reduction in invasion following twice-daily LIV (Fig. 2c).

**Figure 2.**
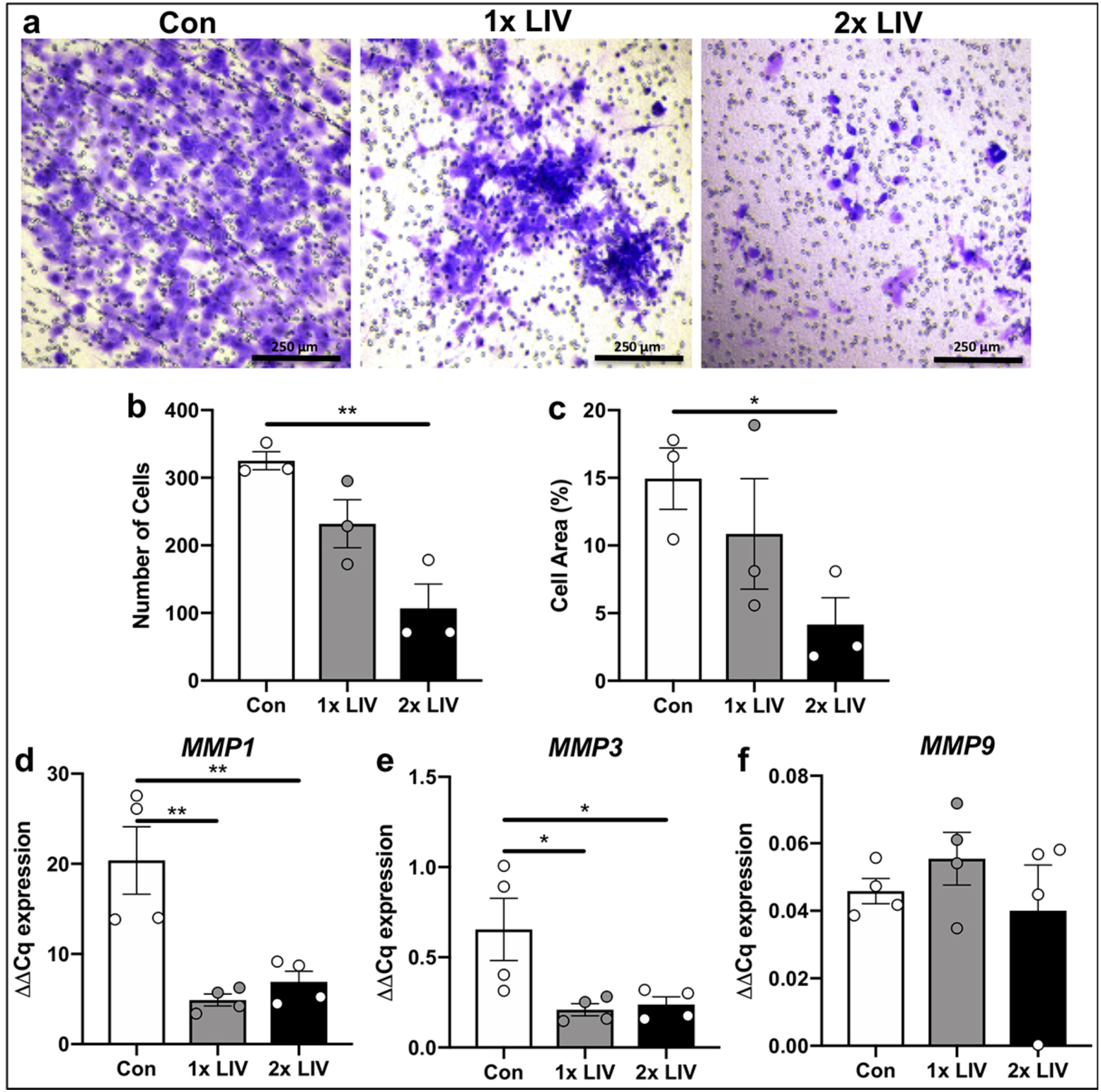
LIV suppresses invasion of MDA-MB-231 cells. (a) Representative images showing crystal violet staining (purple) of MDA-MB-231 cells exposed to once- (1x LIV) or twice-daily (2x LIV) LIV that have invaded through Matrigel^®^ and penetrated through the trans-well membrane. Images are representative of three biological replicates. (b) Quantification of crystal violet stained cells that invaded through the trans-well membrane. Compared to non-vibrated controls (Con) once-daily LIV (1x LIV) induced no significant change, while twice-daily LIV (2x LIV) resulted in a significant reduction in invasion (n=3). (c) Invasion through the trans-well membrane was also quantified by measuring the total area of crystal violet stained cells, relative to the total area of the membrane (n=3). Similar results were found with a significant reduction in invasion following twice-daily LIV (2x LIV). (d) Expression of matrix metalloproteinase 1 (*MMP1*) mRNA (n=4). (e) Expression of *MMP3* mRNA by qPCR (n=4). (f) Expression of *MMP9* mRNA by qPCR (n=4). One-way ANOVA p-values: *p<0.05, **p<0.01.

As invasion through the ECM requires matrix metalloproteinases (MMPs)(Duffy et al., 2000), MMP mRNA levels were quantified by qPCR. *MMP1* expression was decreased by 4-fold (p<0.01) following once-daily LIV and by 3-fold (p<0.01) with twice-daily LIV (Fig. 2d). Expression of *MMP3* was reduced by 3-fold (p<0.05) following once-daily LIV and by 2.7-fold (p<0.05) with twice-daily LIV (Fig. 2e). No significant changes in *MMP9* expression were observed (Fig. 2f).

### Exposure of breast cancer cells to LIV impairs osteoclastogenesis

Breast cancer readily metastasizes to bone, resulting in osteolysis through increased osteoclast formation(Weilbaecher et al., 2011). Tumor-mediated activation of osteoclasts results in an enhanced state of bone resorption and release of matrix-derived growth factors, further stimulating tumor cell invasion and growth(Weilbaecher et al., 2011). LIV restricts cancer-induced bone loss(Pagnotti et al., 2012; Pagnotti et al., 2016), possibly the result of reduced secretion of pro-osteolytic factors from the tumors themselves. To determine if direct application of LIV to cancer cells impairs osteoclastogenesis, MDA-MB-231 cells were exposed to LIV once- or twice-daily, as described. Conditioned media (CM) was collected from MDA-MB-231 cultures 3 hours after the last LIV treatment. Murine RAW 264.7 macrophages were exposed to media from control (non-LIV treated) or LIV treated MDA-MB-231 cells and stained for tartrate resistant acid phosphatase (TRAP). RAW 264.7 cells exposed to media from control MDA-MB-231 cells readily differentiated into multinucleated (>3 nuclei), TRAP positive, osteoclasts; whereas treatment with LIV, once- or twice-daily restricted osteoclast formation (Fig. 3a). Quantification of TRAP stained cultures demonstrated a significant (p<0.0001) reduction in the number of multinucleated cells (≥3 nuclei) following exposure of RAW 264.7 cells to conditioned media from MDA-MB-231 cells vibrated once- (7-fold) or twice-daily (3.5-fold) LIV (Fig. 3b).

**Figure 3.**
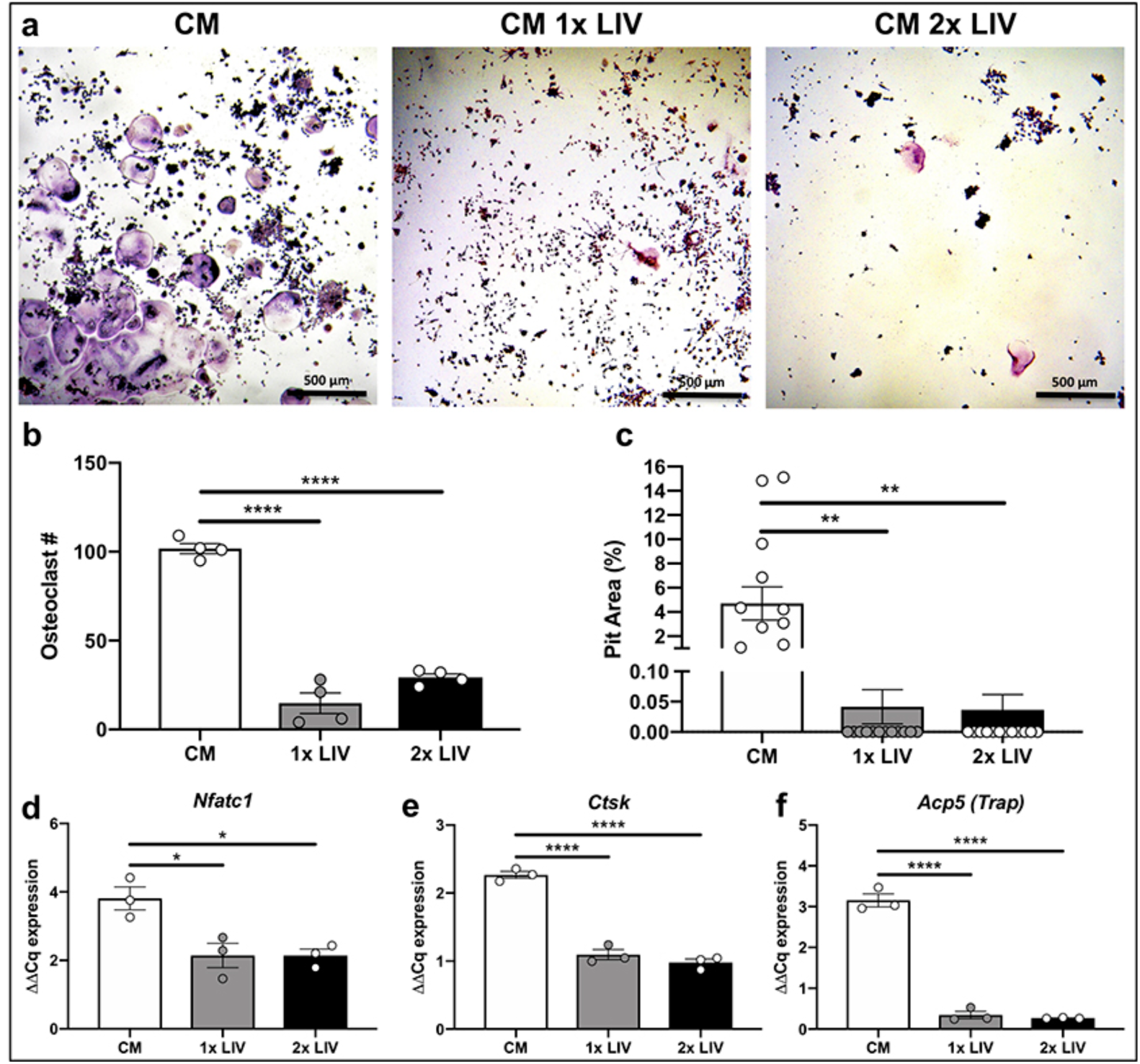
Mechanical stimulation of breast cancer cells releases factors that suppress osteoclastogenesis. (a) Representative images of murine RAW 264.7 cells stained with TRAP after addition of conditioned media from MDA-MB-231 cells not receiving LIV (CM), receiving once-daily LIV (CM 1x LIV) or twice-daily LIV (CM 2x LIV). Images are representative of four biological replicates. (b) Quantification of the number of osteoclasts per area, following treatment with CM from MDA-MB-231 cells. Cells were counted as osteoclasts if they contained positive staining for TRAP (purple) and contained ≥3 nuclei (n=4). (c) RAW 264.7 cells were plated on Osteoassay wells containing hydroxyapatite and exposed to conditioned media from MDA-MB-231 cells, as described above. The area of hydroxyapatite that was resorbed away from the dish was quantified using ImageJ and normalized to the total area (n=10). (d) Quantification of *Nfatc1* mRNA from differentiated RAW 264.7 cells following addition of CM from MDA-MB-231 cells (n=3). Quantification of mRNA from differentiated RAW 264.7 cells following addition of CM from MDA-MB-231 cells for genes that regulate osteoclast differentiation including (d) nuclear factor of activated T-cells (*Nfatc1*), (e) cathepsin-K (*Ctsk*), (f) tartrate resistant acid phosphatase (*Acp5*). Data for each graph represent three biological replicates. One-way ANOVA p-values: *p<0.05, **p<0.01, ****p<0.0001.

To assess the activity of osteoclasts exposed to conditioned media from MDA-MB-231 cells, RAW 264.7 cells were seeded to osteoassay wells (Corning, Corning NY) containing hydroxyapatite. Cells were treated with conditioned media from MDA-MB-231 cells exposed to once- or twice-daily LIV. RAW 264.7 cells exposed to conditioned media from once- and twice-daily LIV treated MDA-MB-231 cells reduced (p<0.01) the resorption pit area by 113-fold and 129-fold respectively (Fig. 3c). These data demonstrate that mechanically stimulating MDA-MB-231 breast cancer cells reduces their ability to support both osteoclast formation and activity.

Osteoclast formation requires the transcription factor nuclear factor of activated T cells 1 (*NFATC1*) and is accompanied by the subsequent production of genes including cathepsin K and *Trap*(Negishi-Koga and Takayanagi, 2009). To determine how conditioned media from LIV treated breast cancer cells reduced osteoclast formation, RNA from RAW 264.7 cells was examined by qPCR following exposure to CM. Exposure of RAW 264.7 cells to CM from MDA-MB-231 cells treated with both once- and twice-daily LIV reduced *Nfatc1* expression by ∼2-fold (p<0.05) for both treatment bouts (Fig. 3d). Expression of cathepsin K (*Ctsk*) was reduced (p<0.01) in cells treated with CM from once- (2-fold) and twice-daily (2-fold) treated MDA-MB-231 cells (Fig. 3e). *Trap* (*Acp5*) expression was also reduced (p<0.05) following treatment with CM from once- and twice-daily LIV treated MDA-MB-231 cells, each with reductions of ∼12-fold (Fig. 3f). These data suggest that the reductions in osteoclast number, following exposure to CM from mechanically stimulated breast cancer cells, result from suppression of osteoclast regulatory genes, including *Nfatc1*, *Ctsk*, and *Acp5*.

### LIV suppresses expression of factors that promote osteolysis

Within the bone microenvironment, cancer cells produce factors that perpetuate osteoclast formation and subsequent osteolysis. As we found that CM from human breast cancer cells, exposed to LIV signals, suppressed osteoclast formation, we next sought to quantify expression of factors produced by MDA-MB-231 cells that promote osteolysis, following LIV. As TGF-β enhances the metastatic phenotype of cancer cells, and increases production of osteolytic factors(Guise and Chirgwin, 2003), MDA-MB-231 were treated with TGF-β1 (5 ng/ml, w/v) or PBS (veh) as a control, prior to LIV. Exposure of MDA-MB-231 cells to LIV for 20 minutes once per day for 3 days resulted in a 2.5-fold decrease of parathyroid hormone like hormone (*PTHLH*) expression (p<0.01) in the absence of TGF-β1 and a decrease of 1.4-fold (p<0.05) in the presence of TGF-β1 (Fig. S1a). There were no changes in expression of connective tissue growth factor (*CTGF*), *CXCR4*, or interleukin 11 (*IL-11*) following once-daily LIV (Fig. S1b, c, d).

Previous work in non-tumor cells showed that twice-daily LIV resulted in enhanced anabolic effects compared to once-daily treatment(Sen et al., 2011), likely due to a “priming” effect following the first bout of mechanical force. To determine if a similar priming effect occurs in breast cancer cell expression of pro-osteolytic/metastatic genes in MDA-MB-231 cells were also quantified following twice-daily LIV treatment for 3 days. Treatment with LIV twice a day resulted in reduced *PTHLH* expression in both the absence (6-fold, p<0.01) and presence (2.6-fold, p<0.05) of TGF-β1 (Fig. 4a). LIV induced a reduction of connective tissue growth factor (*CTGF*) by 2-fold (p<0.05) in the absence of TGF-β1; however, no differences were found in the presence of TGF-β1 (Fig. 4b). Expression of *IL-11* mRNA was reduced both in the presence (∼2-fold, p<0.01) and absence (3-fold, p<0.05) of TGF-β1 following twice-daily LIV. Levels of C-X-C chemokine receptor type 4 (*CXCR4*), interleukin 8 (*CXCL8*), and hypoxia-inducible factor 1-alpha (*HIF1A*) were also quantified and no differences were observed with LIV, regardless of the addition of TGF-β1 (Fig. 4d, e, f).

**Figure 4.**
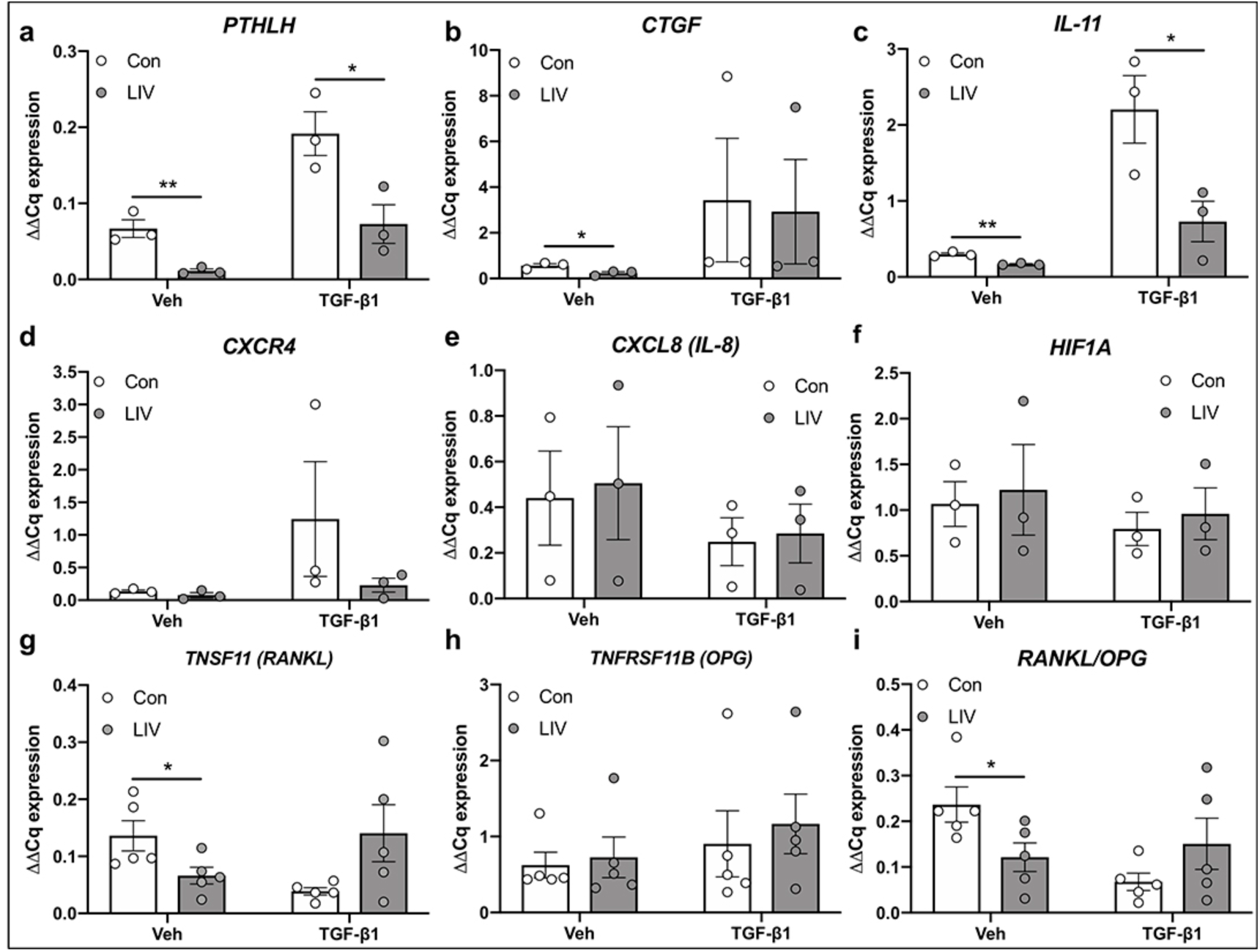
Low-magnitude mechanical forces suppress expression of osteolytic genes. MDA-MB-231 cells were treated with PBS (Veh) or TGF-β1 and exposed to non-vibration control conditions (Con) or LIV twice-daily for 3 days. qPCR analyses were normalized to *GAPDH*. Genes surveyed included (a) parathyroid hormone-related protein (*PTHLH*) (n=3), (b) connective tissue growth factor (*CTGF*) (n=3), (c) interleukin 11 (*IL-11*) (n=3), (d) C-X-C chemokine receptor type 4 (*CXCR4*) (n=3), (e) interleukin 8 (*CXCL8*) (n=3), (f) hypoxia-inducible factor 1-alpha (*HIF1A*) (n=3), (g) receptor activator of nuclear factor kappa-B ligand (TNFSF11, *RANKL*) (n=5), and (h) osteoprotegerin (*TNFRSF11B, OPG*) (n=5). (i) Quantification of the ratio of *RANKL* to *OPG* mRNA (n=5). Multiple t-test p values: *p<0.05, **p<0.01.

Osteoclast formation requires binding of receptor activator of nuclear factor kappa-B ligand (RANKL) to the RANK receptor. While thought to be predominantly produced by osteoblasts, RANKL is also secreted by breast cancer cells(Cross et al., 2006; Sato et al., 2013; Van Poznak et al., 2006). In contrast to RANKL, osteoprotegerin (OPG) suppresses osteoclast formation by acting as a decoy receptor for RANKL(Van Poznak et al., 2006). Exposure of MDA-MB-231 cells to twice-daily LIV reduced *RANKL* expression by 2-fold (p<0.05). When treated with TGF-β1, a non-significant increase of 3.6-fold in RANKL was found following LIV treatment (Fig. 4g). *OPG* expression was not altered with LIV in the presence or absence of TGF-β1 (Fig. 4h); however, the ratio of *RANKL* to *OPG* expression was reduced by 2-fold (p<0.05) in the absence of TGF-β1 and not significantly altered in the presence of TGF-β1 (Fig. 4I). These results demonstrate that LIV suppresses the production of genes in breast cancer cells that influence osteoclast formation.

### Low magnitude mechanical signals increase expression of LINC complex genes

High magnitude mechanical forces, including membrane deformation and fluid shear stress, activate intracellular signaling through focal adhesions (FAs) at the plasma membrane(Thompson et al., 2013). In contrast, low magnitude vibrational forces (<1g) do not induce appreciable fluid shear forces, nor deformation of the plasma membrane(Uzer et al., 2012; Uzer et al., 2013; Uzer et al., 2014), suggesting that low magnitude signals are not transmitted through the cell membrane. Recent work showed that the nucleus acts as a critical mechanosensory organelle and that transmission of low magnitude mechanical signals requires connections between the nucleus and the actin cytoskeleton, which are mediated by the LINC complex(Uzer et al., 2015). To determine if LIV influenced expression of LINC complex genes, Nesprin and Sun, MDA-MB-231 cells were treated with recombinant TGF-β1 or PBS (veh) and subjected to LIV twice daily for three days. Exposure to LIV, in the absence or presence of TGF-β1 resulted in a 1.82-fold (p<0.01) and 1.84-fold (p<0.05) increase in *SYNE1*, the gene encoding Nesprin 1, expression respectively (Fig. 5a). *SYNE2* (Nesprin 2) mRNA was increased by 2.63-fold (p<0.05) in the absence and 1.54-fold (p<0.01) in the presence of TGF-β1 (Fig. 5b). The molecular weight of Nesprin1 (>1,000 kDa) presents challenges for resolving this large protein via Western blotting, thus immunocytochemistry was performed. Cells exposed to 3 days of twice-daily LIV had increased Nesprin1 fluorescence signal compared to non-vibrated controls (Fig. 5c).

**Figure 5.**
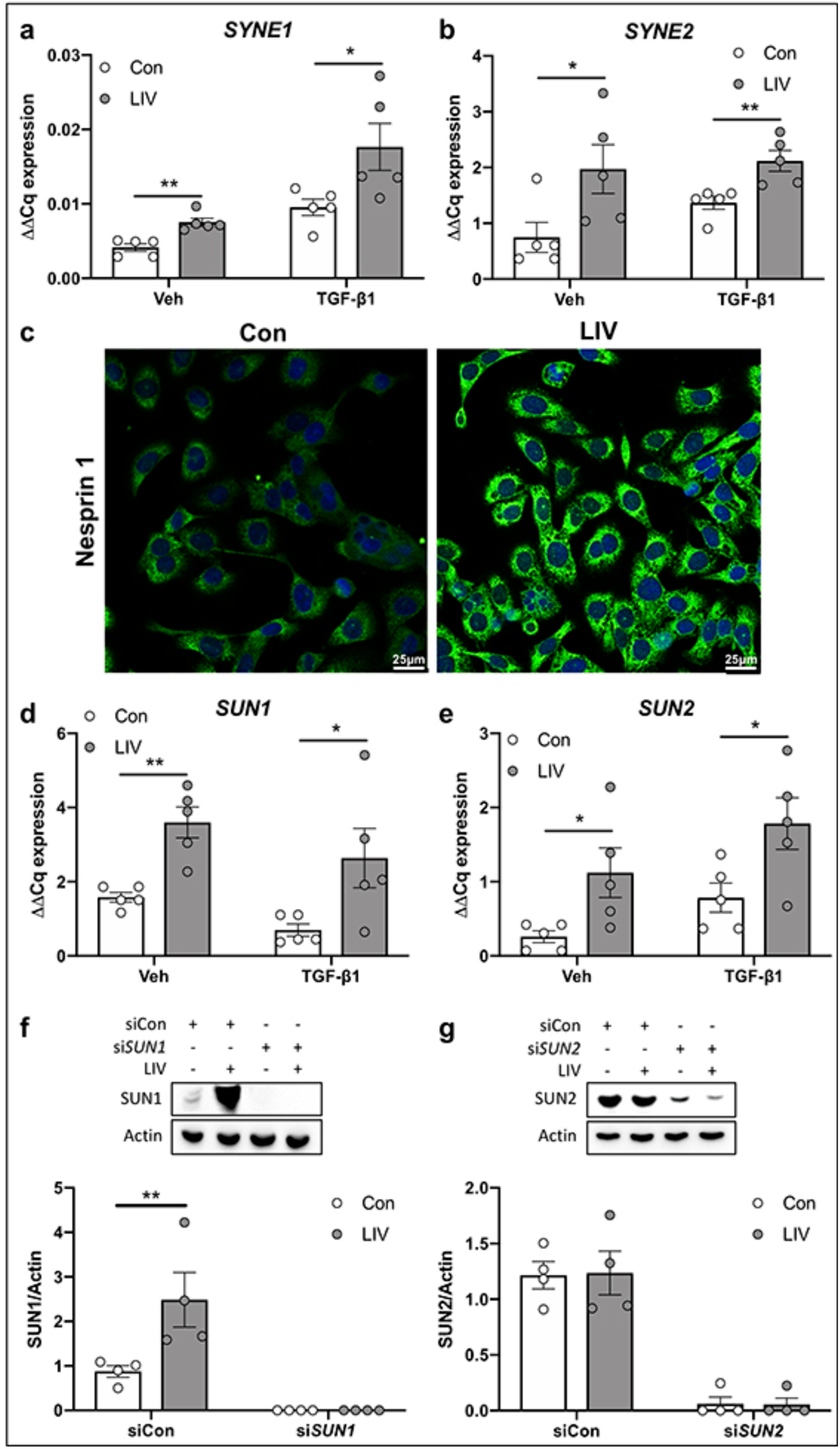
LIV upregulates production of LINC complex components. qPCR analyses of (a) nesprin 1 (*SYNE1*) and (b) nesprin 2 (*SYNE2*) mRNA from MDA-MB-231 cells treated with PBS (Veh) or TGF-β1 and exposed to non-vibration control conditions (Con) or LIV twice-daily for 3 days. qPCR analyses were normalized to *GAPDH* (n=5). (c) Representative images of immunocytochemistry staining of nesprin 1 in MDA-MB-231 cells treated twice-daily with LIV showing increased nesprin 1 signal following LIV. Images are representative of three biological replicates. qPCR analyses of (d) sun 1 (*SUN1*) and (e) sun 2 (*SUN2*) mRNA from MDA-MB-231 cells treated with PBS (Veh) or TGF-β1 and exposed to non-vibration control conditions (Con) or LIV twice-daily for 3 days. qPCR analyses were normalized to *GAPDH* (n=5). Western blots of whole cell lysates (20 μg per lane) from MDA-MB-231 cells transfected with a control siRNA (siCon) or (f) siRNA targeting *SUN1* (si*SUN1*) or (g) siRNA targeting *SUN2* and exposed to non-vibration control conditions (Con) or LIV twice-daily for 3 days. PVDF membranes were blotted with an antibody recognizing SUN1 and β-actin as a loading control. Densitometry was measured using ImageJ and normalized to β-actin. Western blot images are representative of four biological replicates. Uncropped images shown in supplementary figure 5. Multiple t-test p values: *p<0.05, **p<0.01.

Transcript levels of *SUN1* (Fig. 5d) and *SUN2* (Fig. 5e) were increased by ∼2-4 fold following LIV both in the absence and presence of TGFβ. Protein expression of SUN1 and SUN2 was assessed by Western blotting. SUN1 was increased by 2.8-fold (p<0.01) following LIV (Fig. 5f), while SUN2 was not altered (Fig. 5g). Treatment with siRNA targeting *SUN1* (Fig. 5f) or *SUN2* (Fig. 5g) knocked down each protein to nearly undetectable levels. Full size, non-cropped blots are shown in Fig. S5. These findings demonstrate that LIV enhances transcript expression of *SYNE1*, *SYNE2*, *SUN1*, and *SUN2* while increasing protein expression of both Nesprin1 and SUN1 in human breast cancer cells.

### LIV-induced suppression of osteolytic factors requires the LINC complex

The increase in expression of LINC complex genes, following LIV, suggested that low magnitude mechanical forces enhance LINC connectivity, possibly accounting for the downregulation of factors that support osteoclast formation following LIV. As such, MDA-MB-231 cells were transfected with a control siRNA, of similar sequence as the targeting siRNA with variation of critical nucleotides, or siRNA targeting both *SUN1* and *SUN2,* to physically disconnect the LINC complex from the actin cytoskeleton. siRNA targeting both transcripts simultaneously was used as *SUN1* and *SUN2* have redundant functions for binding actin(Lei et al., 2009). Cells were exposed to LIV twice daily for 3 days, as described. While expression of *PTHLH* was significantly reduced (2.5-fold, p<0.01) in cells treated with a control siRNA following LIV, knockdown of *SUN1* and *SUN2* (denoted as *SUN1/2*) resulted in the inability of LIV to suppress *PTHLH* (Fig. 6a). LIV treatment resulted in no changes in expression of *CTGF* with control siRNA or siRNA targeting *SUN1/2* (Fig. 6b). LIV-mediated downregulation of *IL11* expression (1.9-fold, p<0.05) was mitigated following knockdown of *SUN1/2* (Fig. 6c). Expression of *TNSF11* (*RANKL*) was suppressed by LIV with intact *SUN1/2* expression (2.3-fold, p<0.05) and following siRNA-mediated *SUN1/2* knockdown (3.8-fold, p<0.05) (Fig. 6d). LIV had no impact on expression of *TNFRSF11B* (*OPG*) (Fig. 6e); however, when the *RANKL/OPG* ratio was evaluated, the suppression observed with LIV treatment (2.2-fold, p<0.05) was negated following knockdown of *SUN1/2*. These data demonstrate that LIV-mediated downregulation of osteoclastic factors produced by breast cancer cells require an intact LINC complex.

**Figure 6.**
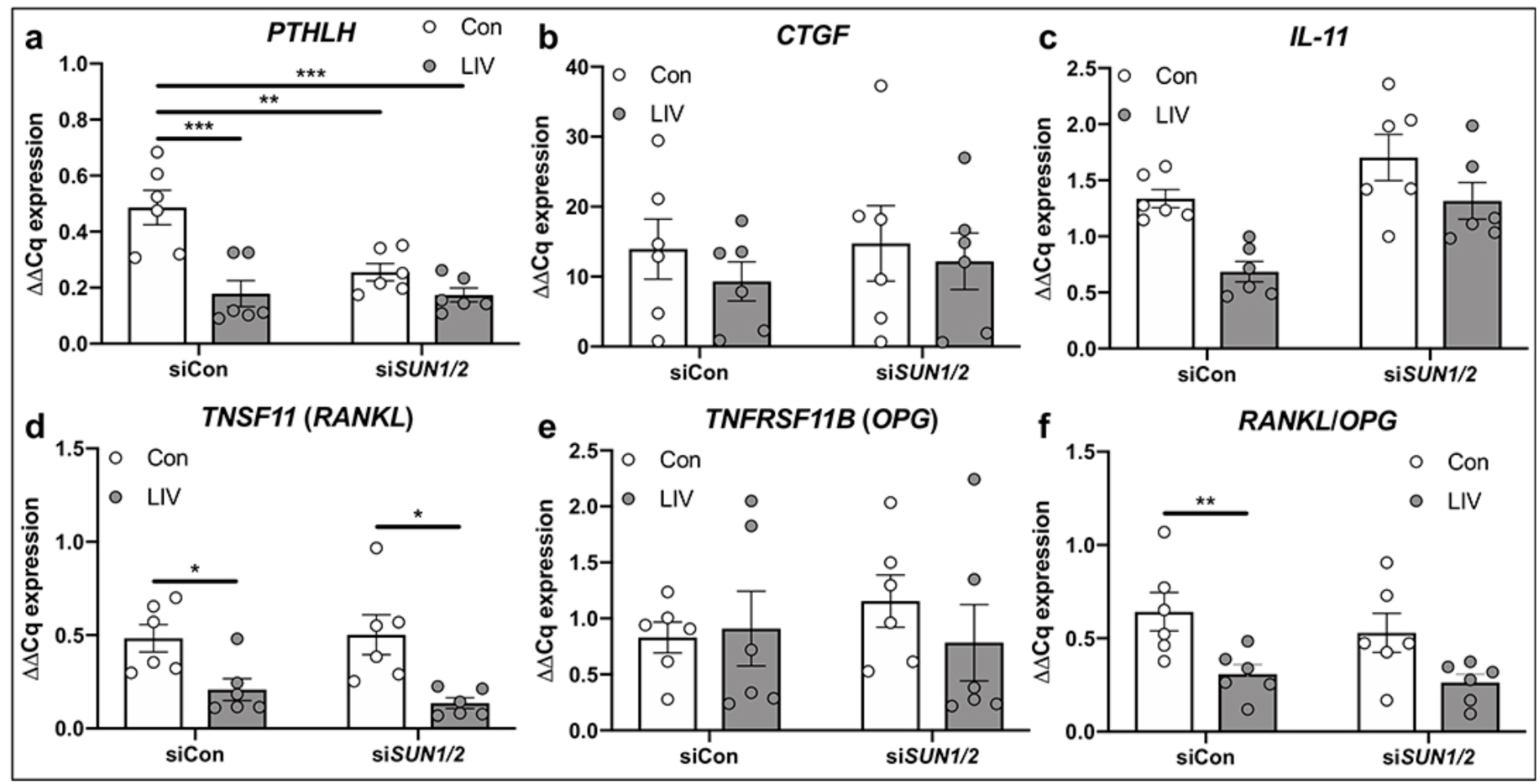
SUN1 and SUN2 are necessary for mechanical repression of osteolytic genes. MDA-MB-231 cells were treated with control siRNA sequences (siCon) or siRNA targeting *SUN1* and *SUN2* and exposed to non-vibration control conditions (Con) or LIV twice-daily for 3 days. qPCR analyses were normalized to *GAPDH*. Genes surveyed included (a) *PTHLH*, (b) *CTGF*, (c) *IL-11*, (d) TNFSF11 (*RANKL*), and (e) *TNFRSF11B* (*OPG*). (i) Quantification of the ratio of *RANKL* to *OPG* mRNA. All graphs represent six biological replicates (n=6). Two-way ANOVA p values: *p<0.05, **p<0.01, ***p<0.001.

### LIV regulates secretion of osteolytic factors from MDA-MB-231 cells

To determine if protein production and secretion of factors that promote osteolysis are influenced by LIV, conditioned media was collected from MDA-MB-231 cells treated with control siRNA or siRNA targeting *SUN1/2* following 3 days of twice-daily LIV. No differences were observed in PTHrP production following LIV with control siRNA or *SUN1/2* siRNA (Fig. S2a). Consistent with transcript levels, secretion of IL11 was reduced following LIV (1.8-fold, p<0.05), while knockdown of *SUN1/2* resulted in increased IL11 production with LIV (2.8-fold, p<0.05) (Fig. S2b). Secretion of RANKL was reduced by LIV (2.6-fold, p<0.5) in cells treated with control siRNA; however, knockdown of *SUN1/2* resulted in a non-significant increase in RANKL (1.9-fold) (Fig. S2c). While there were no changes in OPG secretion with control siRNA or si*SUN1/2* (Fig. S2d), the ratio of RANKL to OPG was decreased in cells transfected with control siRNA (1.7-fold, p<0.05) but increased (2.1-fold, p>0.5) following *SUN1/2* knockdown (Fig. S2e). These data indicate that exposing breast cancer cells to twice-daily LIV reduces secretion of factors that support osteoclastogenesis, and the LINC complex is necessary for these changes.

### LIV-mediated suppression of invasion and osteoclastogenesis requires the LINC complex

To determine if the LINC complex is necessary for the functional effects of LIV on invasion of MDA-MB-231 cells, and the ability of secreted factors to support osteoclast formation, *SUN1/2* were knocked down using siRNA, and cells were treated with LIV as described. The 3-fold reduction in invasion of MDA-MB-231 cells through Matrigel (p<0.05) in cells treated with a control siRNA was negated following knockdown of *SUN1/2*, where no differences in invasion were observed (Fig. 7a & Fig. S3).

**Figure 7.**
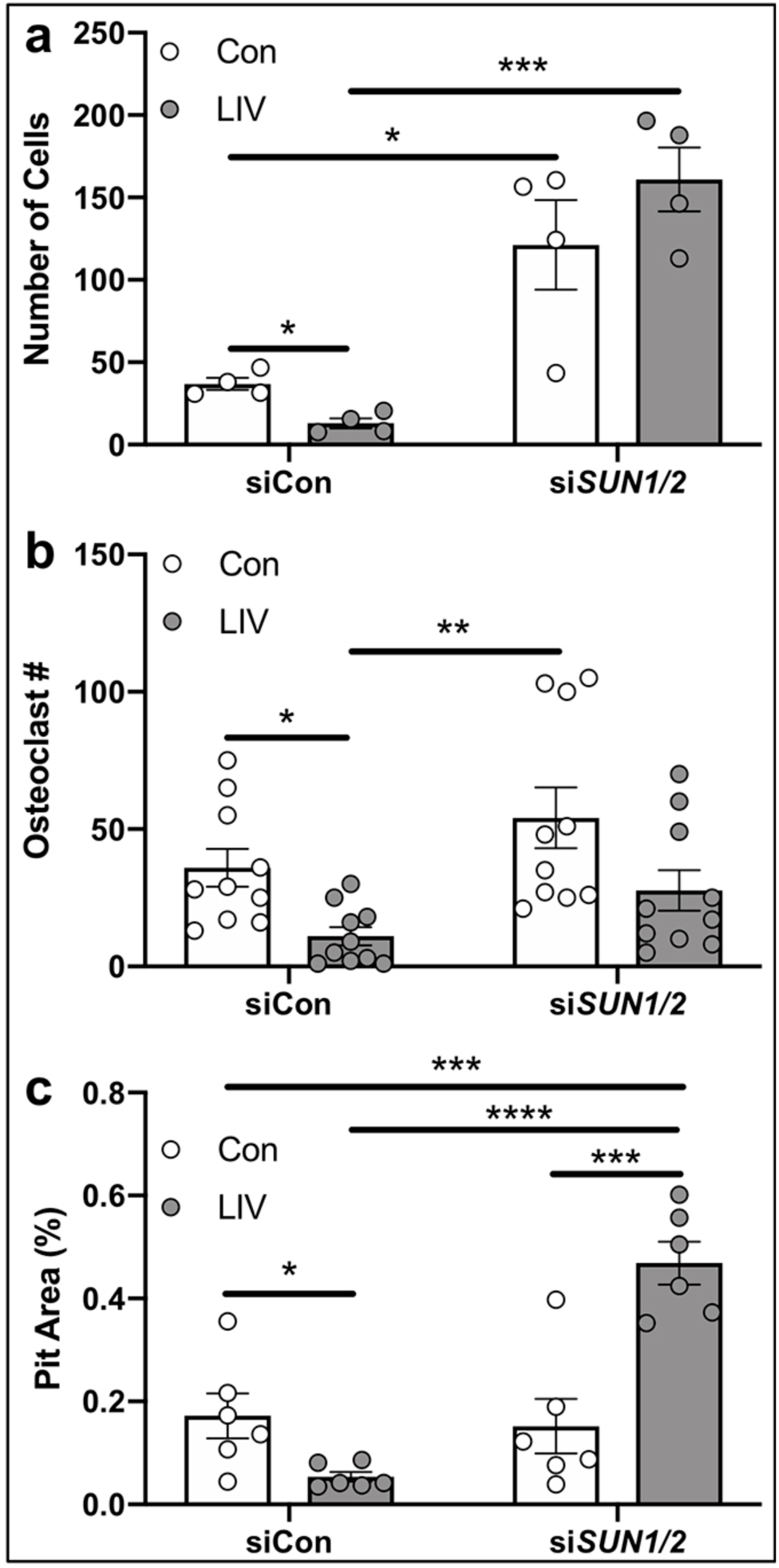
Mechanical suppression of breast cancer cell invasion and signaling to osteoclasts requires SUN1 and SUN2. (a) Quantification of cells (normalized to total area) invading through trans-well membrane under control (Con, no LIV) or twice-daily LIV (LIV) conditions. Cells were transfected with control siRNA oligos (siCon) or with siRNAs targeting both *SUN1* and *SUN2* (si*SUN1/2*). Data is representative of four biological replicates. Representative images are shown in supplementary figure 3 (Fig. S3). (b) Quantification of the number of osteoclasts following exposure of RAW 264.7 cells to conditioned media from MDA-MB-231 cells that received LIV twice-daily for 3 days and transfected with control siRNA sequences (siCon) or siRNAs targeting *SUN1* and *SUN2* (si*SUN1/2*) prior to LIV. Data were compiled from ten biological replicates and representative images are shown in supplementary figure 4 (Fig. S4). (c) Osteoclast resorption, as measured by the area (%) of hydroxyapatite resorbed by differentiated RAW 264.7 cells on Osteoassay plates following the addition of conditioned media from MDA-MB-231 conditioned media. MDA-MB-231 cells were transfected with siRNA sequences targeting *SUN1* and *SUN2* (si*SUN1/2*), as mentioned above, and exposed to twice-daily LIV, or non-LIV control conditions (Con). Data compiled from 6 biological replicates and measured using ImageJ. Two-way ANOVA p values: *p<0.05, **p<0.01, ***p<0.001, ****p<0.0001.

Osteoclast formation of RAW 264.7 macrophages was evaluated following treatment with CM from non-vibrated MDA-MB-231, or from cells exposed to LIV (Fig. S4). CM from LIV treated cells resulted in a reduction in multinucleated (>3 nuclei) osteoclasts (3.2-fold, p<0.05); whereas CM from MDA-MB-231 cells, following knockdown of *SUN1/2,* resulted in no differences in formation of osteoclasts (Fig. 7b).

To assess resorption activity, following exposure to CM from MDA-MB-231 cells, RAW 264.7 macrophages were seeded on Osteo Assay wells. Cells exposed to CM from LIV treated MDA-MB-231 cells, transfected with control siRNA, had a 3.2-fold (p<0.05) decrease in resorption area compared to non-vibrated controls (Fig. 7c). In contrast, addition of CM from cells exposed to LIV, but transfected with siRNA targeting *SUN1 and SUN2,* resulted in a 3.3-fold increase in resorption (p<0.0001). These data demonstrate that Sun1 and Sun2 are essential mediators for the effects of LIV on breast cancer cell invasion and support of osteoclast formation and activity.

### The LINC nuclear complex is required for mechanically-induced cellular stiffness

The mechanical integrity of cancer cells is inversely proportional to metastatic potential(Zhou et al., 2017). We hypothesized that the suppression of invasion and decreases in osteoclastogenesis, observed following LIV, may be the result of increased cell membrane stiffness. To quantify membrane stiffness, MDA-MB-231 cells were exposed to LIV twice-daily for 3 days, or to non-vibrated conditions. Cells were transfected with either control siRNAs or siRNAs targeting *SUN1* and *SUN2*, as described. Cellular stiffness (elastic modulus) was measured by atomic force microscopy, as in Fig. 8a. In cells treated with control siRNA, LIV increased cellular stiffness (1.2-fold, p<0.05), while no differences were found in cells after knockdown of *SUN1/2*.

**Figure 8.**
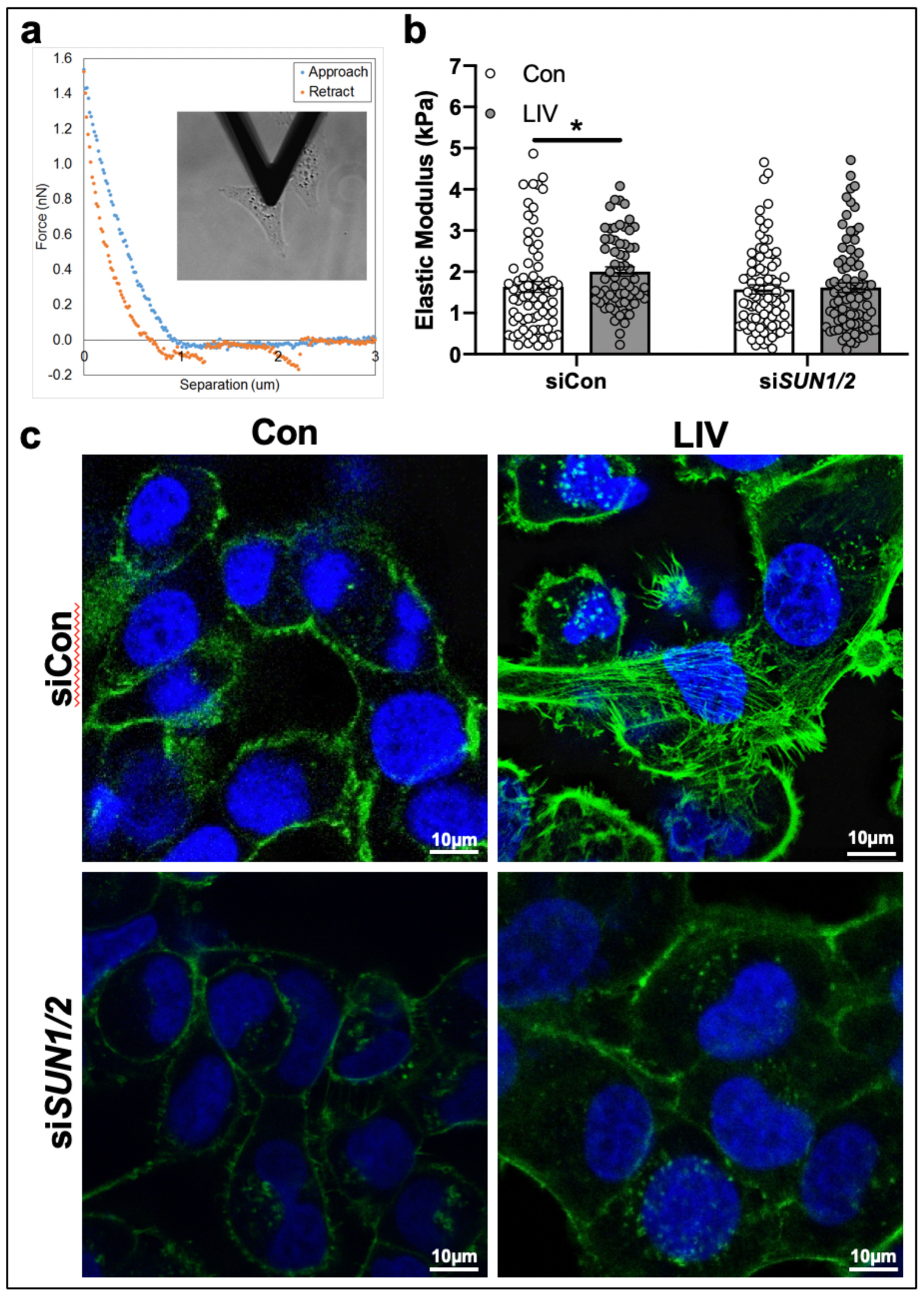
Enhanced cell stiffness, induced by low-magnitude mechanical forces, requires the LINC complex. (a) Image showing the cantilever of the AFM resting over the nucleus of a single MDA-MB-231 cell and representative plot showing forces generated during the approach and retraction of the borosilicate probe (attached to the cantilever) as a function of distance. (b) Graph showing quantification of the elastic modulus (E, kPa). Individual cells from four biological replicate assays were tested. The number of cells tested for each group are as follows… Con/siCon: n=74, LIV/siCon: n=64, Con/si*SUN1/2*: n=85, LIV/si*SUN1/2*: n=85. (c) Representative images of MDA-MB-231 cells exposed to non-vibration control conditions (Con), twice-daily LIV (LIV, and either controls siRNAs or siRNA oligos targeting *SUN1* and *SUN2* (si*SUN1/2*). For each condition, cells were fixed following a 3-hour rest period and incubated with Phalloidin-conjugated Alexa-Fluor 488 and Dapi to stain for filamentous actin (green) and nuclei (blue) respectively. Images are representative of three biological replicates. Statistical analyses calculated using two-way ANOVA, followed by two-stage linear step-up procedure of Benjamini, Krieger, and Yekutieli to control for false discovery rate.

As part of the LINC complex, SUN proteins enable connection between the actin cytoskeleton and the nucleus. Low magnitude mechanical forces enhance actin cytoskeletal structure, which may account for the increased cellular stiffness. Exposure of MB-MD-231 cells to LIV twice daily for 3 days resulted in increased actin cytoskeleton stress fiber formation, as stained by Alexa 488-conjugated phalloidin, compared to non-vibrated controls (Fig. 8c). The increased cytoskeletal structure observed with LIV was negated with knockdown of *SUN1/2*, resulting in phalloidin signal similar to that of non-LIV controls (Fig. 8c).

## Methods

### Reagents

Fetal bovine serum (FBS) was obtained from Atlanta Biologicals (Atlanta, GA). Culture media, trypsin-EDTA, and antibiotics were purchased from Invitrogen (Carlsbad, CA). iTaq universal SYBR green qPCR mastermix (cat# 172-5121) was purchased from Bio-Rad (Hercules, CA). Recombinant human TGFβ-1 (cat# 240B002) was purchased from R&D systems (Minneapolis, MN). Alexa 488-conjugated phalloidin (cat# A12379) was purchased from Invitrogen.

### Cells and Culture Conditions

Human breast cancer cell line, MDA-MB-231, was purchased from the American Tissue Culture Collection (ATCC, Manassas, VA) and cultured in Dulbecco’s Modified Essential Medium (DMEM) supplemented with FBS (10%, v/v) and penicillin/streptomycin (100μg/ml). Murine RAW 264.7 macrophage cells were purchased from ATCC and cultured in DMEM containing FBS (10%, v/v), penicillin/streptomycin (100μg/ml), and recombinant RANKL (75ng/ml). Cells were seeded to densities specific to each protocol.

### Real Time PCR

Total RNA was isolated from cells using RNeasy kit (Qiagen, Germantown, MD), reverse transcribed, and genes were amplified with a BioRad CFX Connect^TM^ qPCR machine, using gene-specific primers, as previously described(Thompson et al., 2015). PCR products were normalized to *GAPDH* and quantified using the ΔΔCT method (denoted as ΔΔCq).

### siRNA-Mediated Knockdown

MDA-MB-231 cells were transfected with gene-specific siRNA or control siRNA (20 nM) using PepMute Plus transfection reagent (SignaGen Labs, Rockville, MD). After 18 hours of transfection, media was replaced with fresh DMEM containing FBS (10%, v/v) and penicillin/streptomycin (100 μg/ml). LIV treatment was initiated 1-2 hours after media was replaced. The following Stealth Select siRNAs (Invitrogen) were used in this study: negative control for *SUN1:* 5’-AAGGTTGCGTGGTTATAAACGCCTG-3’; *SUN1:* 5’-CAGGACGTGTTTAAACCCACGACTT-3’; negative control for *SUN2:* 5’-GCATTACCACCGTCCTTTCGAGGTT-3’; *SUN2:* 5’-GCAGACATTCCACCCTGCTTTGGTT-3’.

### Antibodies

Antibodies targeting SUN1 (cat# ab12) and SUN2 (cat# ab124916) were purchased from Abcam (Cambridge, MA). The Nesprin1 Ab (cat# MA5-18077) was from Invitrogen. The anti-β actin Ab (cat# 5125) was purchased from Cell Signaling (Danvers, MA).

### Low Magnitude Mechanical Force

Low magnitude mechanical forces were applied in the form of LIV using a custom-designed platform, as previously described(Pagnotti et al., 2016). Individual culture dishes were placed on the vertically oscillating platform at room temperature (RT). Cells were stimulated at a frequency of 90 Hz at a magnitude of 0.3 g ± 0.025, where 1 g is equal to the earth’s gravitational field or 9.8 m/s^2^. LIV was applied in 20-minute bouts, once- or twice-daily for 3 days. Twice-daily bouts were separated by 3 hours of rest. Non-vibrated control cells were placed on the vibration platform that was not turned on.

### Cell Viability and Apoptosis Assays

Cell viability was assessed using the CellTiter96® Aqueous One Solution Assay (cat# G3582), which utilizes 3-(4,5-Dimethylthiazol-2-yl)-2,5-Diphenyltetrazolium Bromide (MTT) (Promega, Madison WI). The assay was carried out according to the manufacturer’s protocol. Colorimetric changes were measured at 490nm using a spectrophotometer. Apoptosis was quantified using the BD Pharmingen™ (San Jose, CA) PE Annexin V detection kit (cat# 559763). Cells were stained according to the manufacturer’s protocol and Annexin V labeled cells were quantified using a BD Accuri C6 flow cytometry machine.

### Transwell Assays

Invasion of MDA-MB-231 cells was quantified using Transwell^®^ membrane inserts with a pore size of 8.0 μm (Corning^®^, Corning, NY). Cells first were seeded to 6-well dishes, exposed to LIV or non-LIV conditions for the appropriate number of days. At least 2 hours prior to seeding of cells on to transwell inserts, Matrigel^®^ (Corning^®^, cat# 356234) was diluted with DMEM (1:5) and added (100 μl) to the upper chamber of insert within the 24-well dish. Following LIV, cells were trypsinized and seeded (50,000 cells per well) onto Matrigel^®^-containing upper chamber of the transwell inserts. Cells were suspended in DMEM containing FBS (1%, v/v). The lower chamber contained DMEM with FBS (10%, v/v), P/S (1%, v/v) and collagen (rat tail, type I, cat# 354236, 40 ng/ml). Cells within the transwell insert were incubated at 37°C for 24 hrs, then fixed with methanol (100%, v/v) for 10 min and stained with crystal violet (0.5%, w/v) for 10 min. Cell number and area were quantified under light microscopy.

### Western Blotting

Whole cell lysates were prepared using radio immunoprecipitation assay (RIPA) lysis buffer (150 mM NaCl, 50 mM Tris HCl, 1 mM EGTA, 0.24% sodium deoxycholate,1% Igepal, pH 7.5) containing NaF (25 mM) and Na3VO4 (2 mM). Aprotinin, leupeptin, pepstatin, and phenylmethylsulfonylfluoride (PMSF) were added fresh, just prior to lysis. Whole cell lysates (20 μg) were separated on polyacrylamide gels (4-12%) and transferred to polyvinylidene difluoride (PVDF) membranes. Membranes were blocked with milk (5%, w/v) diluted in TBS-T. Blots then were incubated overnight at 4°C with the appropriate primary antibodies. Blots were washed and incubated with horseradish peroxidase-conjugated secondary antibody (1:5,000 dilution) (Cell Signaling) at RT for one hour. ECL plus was used to detect chemiluminescence (Amersham Biosciences, Piscataway, NJ). Images were developed and acquired with an iBright CL1000 machine (Applied Biosystems), and densitometry was determined using ImageJ software version 1.45s (NIH).

### Immunofluorescence

Following LIV and/or siRNA transfection, cells were fixed with paraformaldehyde (4%, v/v) for 20 min, permeabilized with Triton X-100 (0.1%, v/v) for 5 min at RT, and donkey serum (5%, v/v) blocking buffer diluted in TBS-T was added for 30 min, as previously described(Thompson et al., 2018). Cells were incubated with phalloidin-conjugated Alexa Fluor-488 (Invitrogen) diluted in TBS (1:100) 30 min at RT. Cells were washed, covered, and sealed with mounting medium containing DAPI (Invitrogen).

### Osteoclast Differentiation and Pit Assays

Osteoclast differentiation was determined using murine RAW 264.7 macrophage cells, which were seeded onto 6-well dishes (50,000 cells per well). Cells were maintained in DMEM containing RANKL (75 ng/ml), which was replaced with conditioned media from MDA-MB-231 cells after 24 hrs. A 50% mixture of conditioned media and DMEM was added from each group (+/- LIV and +/- si*SUN1/2*) with RANKL (75 ng/ml). Following 4 days of differentiation, cells were stained with tartrate resistant acid phosphatase (TRAP, Sigma, Cat# 386A-1KT) according to the manufacturer’s protocol. Cells were considered to be osteoclasts if they stained for TRAP and had 3 or more nuclei.

Resorption activity of osteoclasts, in the presence of conditioned media from MDA-MB-231 cells, was quantified using the Osteo Assay (Corning, Corning NY) containing hydroxyapatite. Briefly, 5,000 RAW 264.7 cells were seeded onto the 96-well Osteo Assay plate containing 200 μl of DMEM with RANKL (75 ng/ml). The next day, half the volume of media was replaced with conditioned media from MDA-MB-231 cells for each condition. Media was replaced every two days. At day 4, media was removed and 100 μl of bleach (10%, v/v) was added for 5 min at RT. Bleach was removed and cells were washed twice with water. Formation of pits in the hydroxyapatite were visualized under light microscopy and quantified with ImageJ.

### Determination of Elastic Modulus using Atomic Force Microscopy

Cells were indented in media at RT using a BioScope Catalyst AFM (Bruker, Santa Barbara, CA). The AFM was mounted on a Leica DMI3000 inverted microscope (Leica Biosystems Inc., Buffalo Grove, IL), facilitating accurate placement of the AFM probe over individual cells. Indentations were carried out using a spherical borosilicate bead (5 μm diameter) mounted on a gold-coated silicon nitride cantilever (Novascan Technologies, Inc., Boone, IA). Prior to indenting, probes were pushed onto a glass surface and the deflection of the cantilever was used to measure the cantilever’s deflection sensitivity (nm/V). The cantilever’s spring constant (∼0.07 N/m) was then determined using the thermal tuning method. The light microscope was used to navigate to randomly-selected cells. The apex of the cantilever (where the bead is attached) was placed directly above the nucleus, then the AFM was engaged. A single ramp was performed using a trigger force of 1 nN at a speed of 0.5 Hz.

Analysis was performed using Nanoscope Analysis v1.70 (Bruker). A linear baseline correction was fit from 30-80% of the retraction curve. Since Poisson’s ratio is not fully understood for cells, reduced elastic modulus (E_r_) was fit for each unloading curve in a region spanning from 20-75% of the maximum force using the Hertz model of contact between a rigid sphere and an elastic half space:

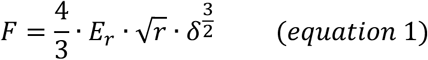

In equation 1, F is force exerted on the cell, r is the radius of curvature of the probe and δ is sample deformation. Only indents with goodness of fits (i.e. r^2^) greater than 0.97 were included for analysis.

### Statistical Analysis

Statistical variance was expressed as the means ± standard error of the mean (SEM). Evaluation of statistical significance performed by one-way analysis of variance (ANOVA), two-way ANOVA, or student’s t-test, as appropriate (Prism GraphPad, La Jolla, CA). For AFM assays, two-way ANOVA was used followed by two-stage linear step-up procedure of Benjamini, Krieger, and Yekutieli to control for false discovery rate. Values were considered significant if p ≤ 0.05. Experiments were replicated at least three times to assure reproducibility. Densitometry data from Western blots were compiled from four biological replicates.

## Discussion

A variety of chemical, hormonal, and physical cues regulate cell behavior; many of which are influenced by exercise. Mechanical signals are a principal component of exercise and are not only instrumental in governing adaptation of musculoskeletal tissues, but also influence cancer cell behavior including invasion(Brabek et al., 2010), division(Provenzano and Keely, 2011), migration(Katira et al., 2012), and metastasis(Wirtz et al., 2011). Here we demonstrate that very low magnitude mechanical signals regulate invasive, apoptotic, and osteoclastogenic properties of human breast cancer cells through biochemical alterations of the cancer cells, via paracrine signaling between tumor cells and osteoclasts, and via biophysical changes in the cancer cells in response to the exogenous mechanical signals. Our data show that mechanical signals directly influence human breast cancer cells by suppressing invasion and increasing the susceptibility to apoptosis. Application of mechanical signals also influenced paracrine signaling, resulting in suppressed osteoclast differentiation and lytic activity. Furthermore, these mechanical signals altered the biophysical properties of cancer cells, resulting in increased membrane stiffness, where force transmission required connection between the nucleus and the actin cytoskeleton, mediated by the LINC complex.

Exercise is an effective form of introducing exogenous mechanical signals to promote musculoskeletal anabolism(Thompson et al., 2016); however, common forms of weight-bearing exercise such as running pose challenges to cancer patients, who are already debilitated due to their illness and subsequent treatments. This can lead to injuries, falls, and fractures, the very thing exercise was intended to prevent. LIV introduces the necessary mechanical signals to maintain musculoskeletal health(Keller et al., 2013; Thompson et al., 2014), yet correct dosing is essential. In non-cancerous cells, delivery of two bouts of mechanical signals separated by a rest period of three hours induced greater anabolic signaling than when the mechanical signals were delivered for the same overall time duration but with no rest period(Sen et al., 2011). These findings mimic responses in mice where incorporation of a refractory period between bouts of mechanical stimulation diminished obesity-induced adipose accumulation(Patel et al., 2017). Our data show that incorporation of a refractory period between bouts of mechanical signals resulted in greater reductions in invasion and expression of osteolytic genes, compared to a single session, suggesting that the initial bout primes the cells, making them more receptive to additional mechanical input. These observations are supported by the increased cytoskeletal structure observed following LIV, where the cells adapt by enhancing actin formation, which activates anabolic signaling cascades(Sen et al., 2014). Furthermore, these data provide important insight for optimal dosing of mechanical signals in clinical settings.

Mechanical signals influence tumor cell death. In one study, fluid shear increased apoptosis in several tumor cell types, while no negative effects of shear were found in non-tumor cells(Lien et al., 2013). In another study, daily application of LIV increased apoptosis of MDA-MB-231 cells(Olcum and Ozcivici, 2014). We did not observe increased cell death directly following LIV; however, LIV increased expression of the membrane death receptor *FAS*, resulting in increased Fas ligand-mediated apoptosis following LIV. As immune cells, including natural killer (NK) cells and T cells(Chua et al., 2004), produce Fas-ligand, our data suggest that LIV renders breast cancer cells more susceptible to immune-mediated apoptosis. In previous work, voluntary running decreased tumor size in mice, an effect that was linked to increased T and NK cell tumor infiltration(Pedersen et al., 2016). Clearing of NK cells from mice negated the effect of exercise(Pedersen et al., 2016). Our data demonstrate that introduction of exogenous mechanical signals regulates susceptibility of immune-mediated responses in breast cancer cells, a possible explanation for altered tumor size following exercise in mice.

In addition to the effects of mechanical signals on cell survival, we found that application of LIV reduced breast cancer cell invasion. These changes were accompanied by reductions in MMPs. As MMPs are associated with increased breast cancer metastases(Bachmeier et al., 2001; Kohrmann et al., 2009), our data suggest that mechanical signals suppress breast cancer metastasis by reducing the ability of tumor cells to invade through matrix.

The skeleton is a frequent site of breast cancer metastasis(Brown and Guise, 2007). Once localized to bone, breast cancer cells release factors, including PTHrP(Guise et al., 1996), IL-11(McCoy et al., 2013), CTGF(Shimo et al., 2006), and RANKL(Canon et al., 2008), which increase osteoclast activity, leading to osteolysis and tumor progression(Esposito et al., 2018; Weilbaecher et al., 2011). As such, we examined the ability of mechanical signals to regulate release of paracrine signals from breast cancer cells. Here we show that direct application of LIV to MDA-MB-231 cells reduced expression and secretion of RANKL and IL-11. Furthermore, when exposed to CM from LIV-treated breast cancer cells, osteoclast precursors had reduced differentiation and resorption activity. Previous work in murine models of ovarian cancer(Pagnotti et al., 2012) and myeloma(Pagnotti et al., 2016) report that LIV diminished the deleterious effects of cancer metastasis on bone structure, specifically by ameliorating cancer-induced osteoclast resorption of trabecular surfaces(Pagnotti et al., 2016). The reduced trabecular erosion may be the result of direct effects of LIV on osteoclasts, or through LIV-mediated alterations in paracrine signaling of cancer cells to osteoclasts. Data supporting the direct effects of LIV on osteoclast differentiation are conflicting(Kulkarni et al., 2013; Sakamoto et al., 2019), where results may be influenced by the regimen of mechanical signals. Our data demonstrate that direct application of LIV to breast cancer cells impairs paracrine signaling necessary for osteoclast activation, an outcome that may be linked to the biophysical responses of the tumor cells.

Mechanical properties of tissues influence behavior. Tumors are stiffer than healthy tissue, due to altered matrix composition(Janmey and Miller, 2011; Mahoney and Csima, 1982). In contrast, the plasma membrane of metastatic cells is less stiff than non-metastatic cells(Cross et al., 2008; Swaminathan et al., 2011; Zhou et al., 2017). Additionally, cell stiffness is increased by enhanced actin cytoskeletal structure, which is associated with decreased metastatic potential(Maeda et al., 2011). Application of mechanical force increases actin cytoskeletal structure(Sen et al., 2014), where increased filamentous actin is frequently interpreted as enhanced cellular stiffness. Using AFM we found that LIV increased the stiffness of human breast cancer cells. This effect was negated following knockdown of *SUN1/2*. These results correlated well with changes in filamentous actin. In contrast to control cells, LIV resulted in an increase of structured actin stress fibers; however, these stress fibers were not seen following knockdown of *SUN1/2* and subsequent LIV. Taken together, these data suggest that application of mechanical stimuli increase cell membrane stiffness of breast cancer cells, possibly accounting for the decreased invasion and release of osteolytic factors. Furthermore, the LINC nuclear complex is necessary to mediate these beneficial effects of LIV, where loss of SUN1 and SUN2 prevent the ability of LIV to influence cell stiffness.

These studies highlight the mechanisms through which mechanical force, a principal component of physical exercise, regulate the invasive potential of cancer cells and their ability to release paracrine signals that influence cancer progression. The role of the nucleus as a mechanosensory apparatus is just beginning to be appreciated, as is the function of the LINC nuclear complex in tumor pathogenicity. However, the use of exogenously applied mechanical forces may be an addition to the therapeutic armamentarium to both reduce tumor progression and prevent musculoskeletal compromise.

## Supporting information

Supplemental figures combined

## Notes

Funding support: This study was supported by Department of Defense BC150678P1 (WRT), NIH AR069943-01 (WRT), NIH AR068332 (US), Department of Defense BC150678 (TAG)

Conflict of Interest: CTR has several issued patents related to the use of mechanical signals to regulate cell behavior, and is a founder of Marodyne Medical, LLC, a developer of LIV technology. All other authors have no conflicts of interest.

