## Supplemental figures combined for "Mechanical Suppression of Breast Cancer Cell Invasion and Paracrine Signaling Requires Nucleo-Cytoskeletal Connectivity"

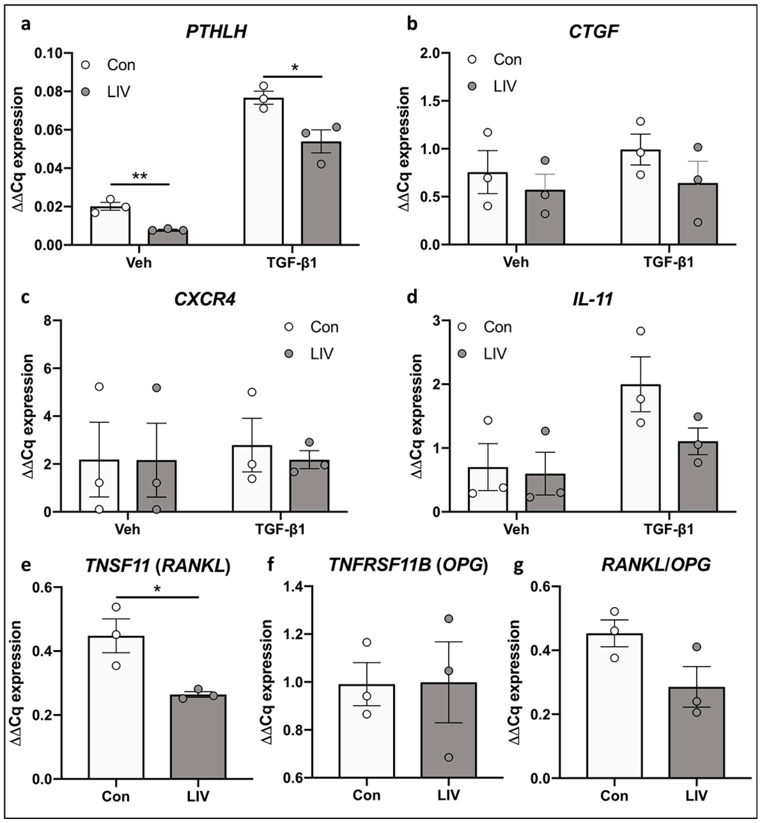


**Figure S1** Expression of osteolytic factors following once-daily LIV. MDA-MB-231 cells were treated with PBS (Veh) or TGF-β1 and exposed to non-vibration control conditions (Con) or LIV once-daily for 3 days. qPCR analyses were normalized to *GAPDH*. Genes surveyed included (a) parathyroid hormone-related protein (*PTHLH*) (n=3), (b) connective tissue growth factor (*CTGF*) (n=3), (c) C-X-C chemokine receptor type 4 (*CXCR4*) (n=3), (d) interleukin 11 (*IL-11*) (n=3), (e) receptor activator of nuclear factor kappa-B ligand (TNFSF11, *RANKL*) (n=3), and (f) osteoprotegerin (*TNFRSF11B, OPG*) (n=3). (g) Quantification of the ratio of *RANKL* to *OPG* mRNA (n=3). Multiple t-test or student's t-test p values: *p<0.05, **p<0.01.

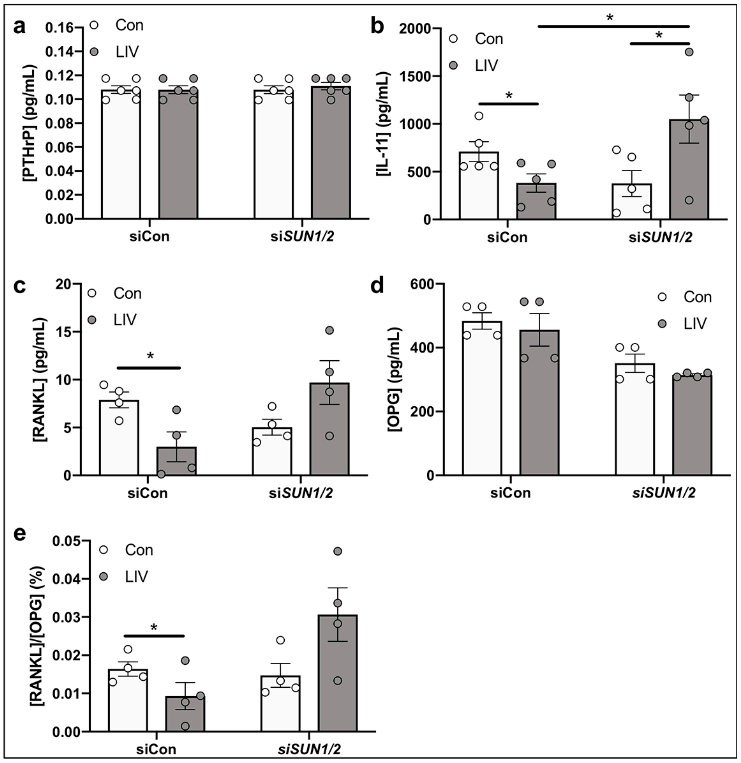


**Figure S2** Secretion of osteolytic factors into conditioned media. Conditioned media was collected from MDA-MB-231 cells following transfection with control siRNAs (siCon) or siRNAs targeting *SUN1* and *SUN2* (siSUN1/2) and non-vibration control conditions (Con) or twice-daily LIV for 3 days (LIV). ELISA assays were performed for the following proteins (a) PTHrP (n=6), (b) IL-11 (n=5), (c) RANKL (n=4), and (d) OPG (n=4). (e) The ratio of RANKL to OPG protein was determined. Multiple t-test or student p values: *p<0.05.

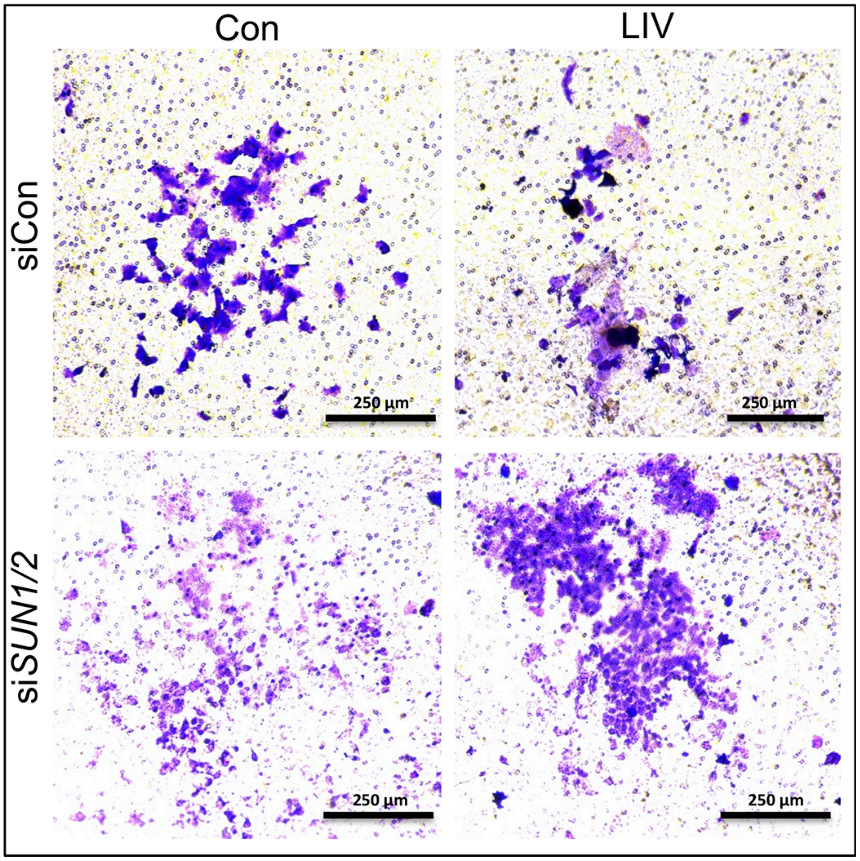


**Figure S3** Mechanical suppression of MDA-MB-231 invasion requires *SUN1* and *SUN2*. Representative images showing crystal violet staining (purple) of MDA-MB-231 cells that have invaded through Matrigel® and penetrated through the trans-well membrane. Cells were exposed to twice-daily LIV for 3 days (LIV) after transfection with control siRNA oligos or siRNAs targeting *SUN1* and *SUN2*. Images are representative of four biological replicates. Quantification of cell invasion provided in figure 7a.

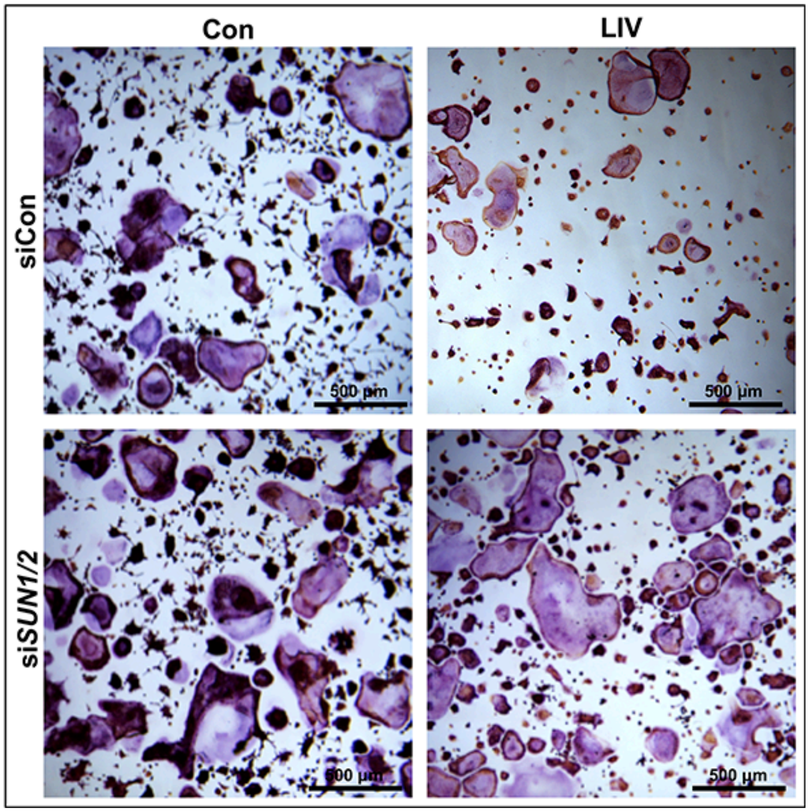


**Figure S4** The LINC complex regulates the ability of mechanical force to alter secretion of factors from breast cancer cells that influence osteoclast formation. Representative images showing RAW 264.7 cells stained with TRAP following the addition of conditioned media from MDA-MB-231 cells that were transfected with control siRNAs or siRNAs targeting *SUN1* and *SUN2* and exposed to non-vibration control conditions (Con) or twice-daily LIV for 3 days (LIV). A 50:50 mixture of conditioned media and growth media (DMEM) was added to each well of RAW 264.7 cells for 4 days prior to TRAP staining. Quantification of osteoclast number, following TRAP staining, is shown in figure 7b.

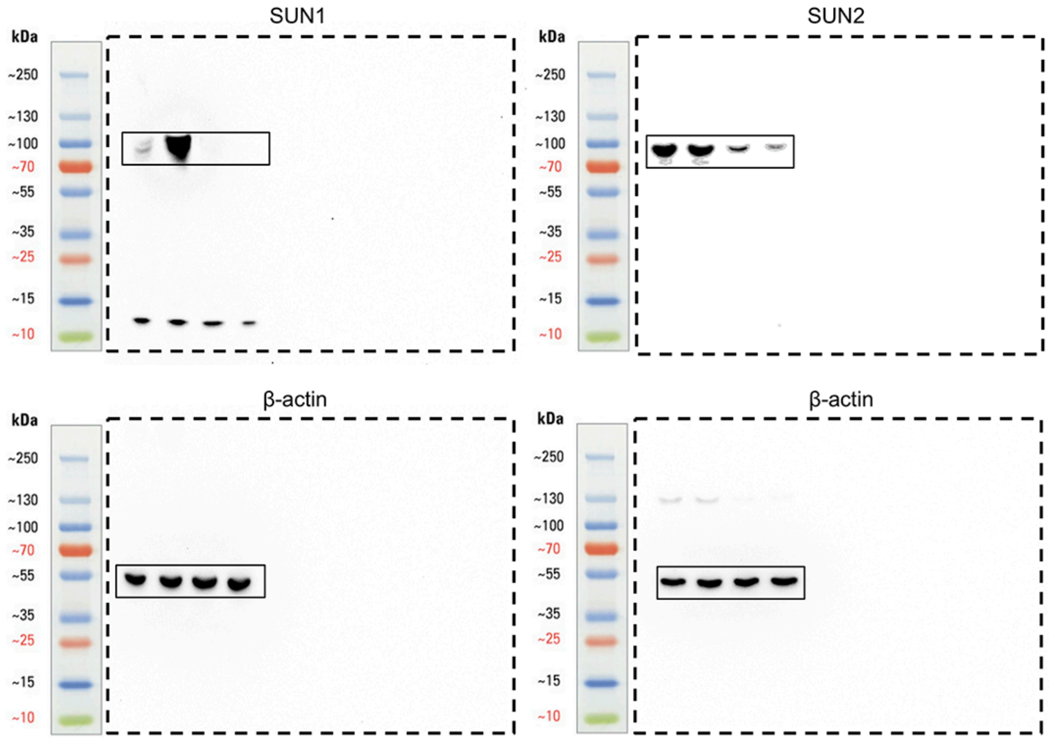


**Figure S5** Original un-cropped images of Western blots shown in figures 6f and 6g. Lysates were collected using RIPA buffer following siRNA knockdown of *SUN1* or *SUN2* (or using control siRNAs) or exposure to twice-daily LIV (or non-LIV conditions), as shown in figure 6.
